# A guide for establishing patient-derived organoids from bile samples obtained during endoscopic procedures and performing gene expression knockdown

**DOI:** 10.64898/2026.03.03.709312

**Authors:** Carla Rojo, Juan J Vila, Laura Guembe, Amaia Arrubla-Gamboa, Vanesa Jusué-Irurita, Juan Carrascosa-Gil, María Rullan, Javier Randez, Maite G Fernández-Barrena, Meritxell Huch, Jesús Urman, Matías A Ávila, Carmen Berasain, Maria Arechederra

## Abstract

Bile represents a clinically accessible biological fluid that can mitigates major limitations associated with tissue-based sampling for the generation of organoid models to study hepatobiliary disease, including biliary tract cancers where tissue availability is often limited. Importantly, bile can also enable the generation of non-malignant cholangiocyte organoids that are otherwise difficult to obtain. Here, we describe an operator-oriented, step-by-step protocol to generate organoids from fresh bile collected during endoscopic retrograde cholangiopancreatography (ERCP), together with two complementary workflows for siRNA delivery in 3D cultures. We detail critical control points that are often under-reported, yet considerably influence success and reproducibility. The protocol was optimized and applied in a real-world cohort of 21 patients undergoing ERCP, including benign biliary obstruction due to choledocholithiasis (n=5) and malignant strictures (n=16: cholangiocarcinoma n=13, gallbladder adenocarcinoma n=1, ampullary tumors n=2). Expandable organoids were established in 17/21 cases (81%), with establishment rates of 60% for choledocholithiasis and 85–100% across malignant entities. Anticipated results include organoid outgrowth within ∼2–3 weeks and morphological heterogeneity in cultures derived from malignant strictures, where normal-like and tumor-like populations may initially coexist and can drift toward a cystic phenotype under routine expansion, motivating optional manual handpicking when tumor-enriched lines are required. As downstream readouts, we show feasibility of DNA-based profiling in selected paired bile–organoid samples (targeted sequencing and ULP-WGS copy-number analysis) and demonstrate proof-of-concept gene silencing via siRNA in both dissociated cells prior to re-embedding, and intact fully formed organoids while preserving 3D architecture. Collectively, this workflow provides a practical and reproducible framework to establish, expand, characterize and functionally perturb bile-derived organoids from routine clinical procedures, facilitating standardized implementation across laboratories.

## 1 Introduction

Organoids are a versatile in vitro platform with broad applications ranging from pathophysiology studies to drug discovery and toxicity testing (Method of the Year 2017: Organoids, 2018; Corrò et al., 2020; Driehuis et al., 2020). Although outcomes may vary across laboratories due to differences in source material, culture conditions, and operator-dependent steps, patient-derived organoids have consistently been shown to retain key genetic and phenotypic features of the original tissue and to provide reproducible, clinically informative experimental models (van de Wetering et al., 2015; Clevers, 2016; Hu et al., 2018; Sachs et al., 2018; Driehuis et al., 2020).

Biliary tract cancers (BTCs), encompassing cholangiocarcinoma (CCA) and gallbladder carcinoma (GBC), are highly lethal malignancies characterized by late diagnosis and limited therapeutic options (Benavides et al., 2015; Banales et al., 2020; Okumura et al., 2021). A major hurdle in BTC research and precision medicine is the scarcity of robust preclinical models that faithfully recapitulate patient’s tumor biology (Amato et al., 2020; Krendl et al., 2025).

Liver organoids can be generated from different sources such as tissue biopsies and pluripotent stem cells (Huch et al., 2013; Takebe et al., 2013; Broutier et al., 2017; Nuciforo and Heim, 2021; Bregante et al., 2026). However, access to the biliary epithelium is often limited, and obtaining non-malignant cholangiocytes as controls is particularly challenging due to the invasiveness of sampling and the suboptimal quality of advanced disease-stage tissues. In this context, bile represents an accessible biospecimen and a practical starting material for biliary organoid derivation (Soroka et al., 2019b; Roos et al., 2021b; Kinoshita et al., 2023). In this context, our group established the Bilebank project (https://bilebank.org/) as an international initiative to promote and advance the use of bile collected during endoscopic retrograde cholangiopancreatography (ERCP) as a liquid biopsy matrix. ERCP is routinely performed in tertiary referral hospitals for both diagnostic evaluation and therapeutic management of biliary strictures originated from benign conditions such as choledocholithiasis, as well as from biliary tract and pancreatic malignancies.

As previously demonstrated, bile collected during endoscopic procedures without representing any additional risk for the patients can be leveraged as a liquid biopsy matrix. For instance, we showed that targeted mutational analysis of bile cell-free DNA can discriminate benign from malignant biliary strictures, supporting early cancer detection (Arechederra et al., 2022, 2025). Moreover, we have shown that protein and lipid-related signatures (Urman et al., 2020), cfDNA methylation biomarkers (Loi et al., 2022), and microbially amidated bile acids (Temprano et al., 2025), can provide complementary clinically relevant information (Arechederra et al., 2024).

Building on the availability of fresh bile samples across benign and malignant biliary diseases, we have optimized a robust workflow to establish bile-derived organoids. In addition, to enable functional interrogation beyond descriptive characterization, we implement an optimized approach for siRNA delivery into 3D cultures, both disaggregated and fully formed, facilitating gene-silencing studies in bile-derived organoids.

## 2 Materials and equipment

### 2.1. Study design

Bile samples were collected from a cohort of 21 individuals with biliary strictures scheduled to undergo endoscopic retrograde cholangiopancreatography (ERCP) at Navarra University Hospital (Pamplona, Spain) between January 2023 and December 2025. Demographic and clinical characteristics are summarized in **Table 1**. The study was approved by the Research Ethics Committee of the Navarra University Hospital (Pamplona, Spain; protocol number CEIm PI_2016/91). All participants were ≥18 years old and provided written informed consent for sample collection, analysis and use of clinical data.

**Table 1.** Characteristics of patients with biliary strictures. Values are median (range) or n (%). Clinical data correspond to patients with biliary strictures undergoing ERCP for bile sampling and organoid derivation. CA19-9 levels were dichotomized according to a clinically relevant cut-off (500 U/mL). Abbreviations: CA19-9, carbohydrate antigen 19-9; ERCP, endoscopic retrograde cholangiopancreatography; nd, not determined.

|  |  | N° | Percentage |
| --- | --- | --- | --- |
| **Gender, n (%)** | Female | 11 | (52.4%) |
|  | Male | 10 | (47.6%) |
| **Median age years, n (range)** |  | 75 | (50-92) |
| **Location of biliary strictures, n (%)** | Distal | 8 | (38.1%) |
|  | Perihilar | 8 | (38.1%) |
|  | nd | 5 | (23.8%) |
| **CA19-9, U/mL n (%)** | <500 | 11 | (52.4%) |
|  | ≥500 | 6 | (28.6%) |
|  | nd | 4 | (19.0%) |

#### 2.1. Human bile samples: collection and transport

Bile was collected from overnight-fasted participants during standard ERCP performed by experienced endoscopists, as previously described (Urman et al., 2020). Briefly, after cannulation of the bile duct, and prior to contrast injection, bile was aspirated through the sphincterotome into sterile additive-free tubes. Usually among 4-15 ml of bile are collected. Samples were immediately transported to the laboratory at 4°C (on ice) and processed within 2 h of collection to initiate the organoid establishment protocol.

**NOTE#1.** Longer pre-processing times were not evaluated in this study.

### 2.2. Reagents and consumables

Reagents for erythrocyte lysis: ACK buffer (ammonium chloride–potassium bicarbonate–EDTA) (prepared in-house; see Section 2.5)

- Ammonium chloride (NH₄Cl) (Sigma-Aldrich; cat. no. A9434)
- Potassium bicarbonate (KHCO₃) (Sigma-Aldrich; cat. no. 60339)
- Disodium EDTA (Sigma-Aldrich; cat. no. E9884)

#### Reagents for organoid culture, passaging and cryopreservation

- Advanced DMEM/F12 (Gibco; cat. no. 12634-010)
- Glutamine/penicillin/streptomycin (100×) (Gibco; cat. no. 10378-016)
- HEPES solution 1 M (Sigma; cat. no. H3537-100ML)
- 3dGRO™ L-WNT Conditioned Media Supplement (Merck Millipore; cat. no. SCM105).
- Primocin, 500x (Invivogen; cat. no. ant-pm-2)
- B27 supplement, 50x, minus vitamin A (Gibco; cat. no. 12587-010)
- N2 supplement, 100x (Gibco; cat. no. 17502-048)
- Nicotinamide (Merk; cat. no. N0636-100G)
- N-Acetylcysteine (NAC) (Sigma-Aldrich; cat. no. A8199-10G)
- Y-inhibitor (Tocris; cat. no. Y-27632)
- [Leu15]-Gastrin I human (Sigma-Aldrich; cat. no. G9145-.1MG)
- Human HGF (Peprotech; cat. no. 100-39H)
- Human FGF10 (Peprotech; cat. no. 100-26)
- Human EGF (Peprotech; cat. no. AF-100-15)
- A83-10 (TGFb inhibitor) (Merck; cat. no. SML0788).
- Forskolin (Calbiochem; cat. no. 344270)
- Dulbecco’S Phosphate Buffered Saline (PBS) (Gibco; cat. no. 14190-094)
- Dimethyl sulfoxide (DMSO) (Sigma Aldrich; cat. no. D2650)
- Bovine serum albumin (BSA) (Sigma Aldrich; cat. no. A9647)
- TrypLE Express Enzyme (Gibco; cat. no. 11598846)
- Recovery™ Cell Culture Freezing Medium (Gibco; cat. no. 12648-010)
- Matrigel Basement Membrane Matrix, LDEV-free, 10 mL (Corning; cat. no. 354234).

**CRITICAL HANDLING**: **1)** Aliquot upon receipt and store at −20 °C (or according to the manufacturer’s recommendations) to avoid repeated freeze–thaw cycles. **2)** Thaw on ice or at 4 °C (e.g., overnight in a 4 °C refrigerator) and never thaw at room temperature. **3)** Maintain Matrigel at 4 °C or on ice during manipulation to prevent premature polymerization.

#### Reagents for organoid Immunohistochemistry

- Agarose (Pronadisa; cat. no. 8016)
- Cryomolds (Bio Optica; cat. no. 07-MP7070)
- Single edge razor blades (Trodis; cat. no. 26025)
- Tissue cassettes (VWR (Avantor); cat. no. 720-2186)
- Absolute ethanol (Panreac; cat. no. 141086.1211)
- 96% ethanol (Oppac; cat. no. 033TC0037)
- Xylene (Prolabo; cat. no. 28973.363)
- Microtome blades (Astelab; cat. no. MCUT-3508BOX)
- TOMO slides (Matsunami; cat. no. TOM-1190)
- Diamond pencil (RS-PRO; cat.no. 394-217)
- Trizma Base (Sigma; cat. no. T6066)
- EDTA (Panreac; cat. no. 131669)
- H_2_O_2_ (Panreac; cat. no. 141077.1211)
- NaCl (Panreac; cat. no. 141659.1211)
- 37% HCl (Panreac; cat. no. 141020.1611)
- Tween® 20 (Sigma; cat. no. P1379)
- Anti-EpCAM (Abcam; cat. no. ab223582)
- Anti-CK19 (Abcam; cat. no. ab52625)
- Envision anti-Rabbit (Dako; cat. no. K4003)
- DAB (+) (Dako; cat. no. K3468)
- Plastic staining jar (Deltalab; cat. no. 19335)
- Glass staining jar (VWR; cat. no. 6319311)

#### Reagents for organoid gene silencing

- Lipofectamine RNAiMAX (Invitrogen; cat. no. 13778075)
- Opti-MEM (Gibco; cat. no. 31985047)
- FBS (Gibco; cat. no A5256701)
- siRNAs (Sigma-Aldrich)

#### Reagents for protein extraction, Western blots, DNA extraction and NGS analysis

Standard reagents and protocols for cell analysis are used.

### 2.3. Equipment

- 24-wells cell culture plates (Nunc™ Thermo Scientific; cat. no. 144530)
- Cryogenic vial, 1 mL (e.g. Thermo Scientific; cat. no. 377267)
- Pipettes P2, P20, P200 and P1000 (e.g. Thermo Fisher)
- Filtered tips (e.g. Thermo Fisher P20 (cat. no. 10731194), P200 (cat. no. 11782584), P1000 (cat. no. 11749855))
- Gel-loading pipet tips (e.g. DeltaLab; cat. no. 200079R)
- Falcon 70 µm nylon cell strainer (Corning/Falcon; cat. no. 352350)
- Class II biological safety cabinet (e.g. TELSTAR; cat. no. BIO-II-A)
- CO₂ cell culture incubator (37°C, humidified, 5% CO₂) (e.g. FORMA SCIENTIFIC; cat. no. 3121)
- Refrigerated centrifuge capable of 200–1800 × g at 4°C (with adaptors for 1.5 mL and 15/50 mL tubes)
- 37°C water bath
- Tube rotator and/or rocker platform (required for steps performed under gentle agitation; e.g., 32°C rocking during siRNA incubation)
- Inverted brightfield microscope for organoid monitoring (e.g., Zeiss; Cell Observer Z1) with a camera for the image acquisition (e.g., Zeiss; Axiocam MRm).
- Rotary microtome (e.g. Microm; cat. no. HM 340E)
- Tissue flotation bath (e.g. MEDITE Medical GmbH; cat. no. TFB 35)
- Pascal pressure chamber (e.g Dako; cat. no. S2800)
- Drying oven (e.g. Indelab; cat. no. IDL-AL-36)
- A digital slide scanner (e.g., Aperio CS2 Digital Scanner, Leica Biosystems) is required to scan the immunohistochemistry preparations.

### 2.4. Reagent setup

#### Reagents for organoid culture mediums

All reagents were prepared under sterile conditions and according to the manufacturers’ recommendations. Stock solutions were prepared at the concentrations indicated below, aliquoted to avoid repeated freeze–thaw cycles, and stored at the appropriate temperatures until use.

- **Nicotinamide (1M):** 250 mg nicotinamide were dissolved in 2.05 mL sterile distilled water, mixed until fully dissolved, aliquoted, and stored at −80 °C
- **N-acetylcysteine (NAC, 500 mM):** 250 mg of NAC were dissolved in 3.06 mL sterile distilled water, aliquoted, and stored at −20 °C for a maximum of 1 month.
- **Y-27632 (ROCK inhibitor, 10 mM):** 2 mg Y-27632 were dissolved in 624 µL sterile PBS, aliquoted, and stored at −80 °C.
- **[Leu15]-Gastrin I human (100 µM):** 0.1 mg peptide were dissolved in 480.7 µL PBS supplemented with BSA (PBS/BSA), aliquoted, and stored at −80 °C.
- **Recombinant human HGF (25 µg/mL):** 25 µg HGF were resuspended in 1 mL PBS/BSA, aliquoted, and stored at −80 °C.
- **Recombinant human FGF10 (100 µg/mL):** 25 µg FGF10 were dissolved in 250 µL PBS/BSA, aliquoted, and stored at −80 °C.
- **Recombinant human EGF (50 µg/mL):** 100 µg EGF were resuspended in 2 mL PBS/BSA, aliquoted, and stored at −80 °C.
- **A83-10 (TGF-β inhibitor, 500 µM):** 0.5 mg A83-10 were dissolved in 2.37 mL DMSO, aliquoted, and stored at −20 °C.
- **Forskolin (8 mM):** 10 mg forskolin were dissolved in 3.045 mL DMSO, aliquoted, and stored at −20 °C.

**Basal culture medium (BCM).** BCM consisted of Advanced DMEM/F-12 (500 mL) supplemented with 5 mL Glutamine/Penicillin/Streptomycin (100× stock; final concentration 1×) and 5 mL HEPES (1 M stock; final concentration 10 mM). The medium was gently mixed and stored at 4 °C until use. BCM was used for washing steps and as the base medium for preparation of complete culture medium (CCM).

**Complete culture medium (CCM)** was prepared by supplementing BCM with Primocin (100 µg/mL), 3dGRO™ L-WNT conditioned medium (1×), N2 supplement (1×), B27 supplement without vitamin A (1×), nicotinamide (10 mM), NAC (1.25 mM), Y-27632 (10 µM), [Leu15]-Gastrin I (10 nM), recombinant human HGF (25 ng/mL), recombinant human FGF10 (100 ng/mL), recombinant human EGF (50 ng/mL), A83-01 (5 µM) and forskolin (10 µM). The final composition and the volumes required to prepare 10 mL CCM are detailed in **Supplementary Table S1**. CCM was gently mixed by inversion and used immediately or stored at 4 °C until use. CCM was prepared weekly. For use, only the required volume was pre-warmed to 37 °C; pre-warmed medium was not returned to 4 °C storage and any leftover volume was discarded.

**10× ACK lysis buffer (1.5 M NH₄Cl, 100 mM KHCO₃, 1 mM EDTA; pH 7.2–7.4; 500 mL).** A 10× ACK lysis buffer stock solution was prepared by dissolving 43 g of NH₄Cl, 5 g of KHCO₃, and 0.186 g of disodium EDTA (EDTA-2Na) in approximately 400 mL of sterile distilled water with stirring until complete dissolution. The pH was adjusted to 7.2–7.4, and sterile distilled water was then added to a final volume of 500 mL. The solution was filter-sterilized through a 0.22 µm filter and stored at 4 °C.

**1× ACK lysis buffer (0.15 M NH₄Cl, 10 mM KHCO₃, 0.1 mM EDTA; pH 7.2–7.4).** A 1× ACK working solution was prepared immediately before use by diluting the 10× stock 1:10 in sterile distilled water.

**2% agarose (w/v) in PBS.** A 2% agarose solution was prepared by weighing 2 g of agarose and adding it to 100 mL of PBS. The suspension was heated in a microwave in several short intervals until boiling and complete dissolution. The solution was then allowed to cool to 55–60 °C in a water bath before use.

**Tris-EDTA buffer (10 mM Tris base, 1 mM EDTA, pH 9.0; 1 L).** A Tris-EDTA (TE) buffer was prepared by dissolving 1.21 g of Trizma Base and 0.37 g of EDTA in 990 mL of deionized water with stirring until complete dissolution. The pH was checked and adjusted to 9.0 if necessary. Deionized water was then added to a final volume of 1 L. The buffer was stored at room temperature for up to 3 months.

**5× Tris Buffered Saline (5× TBS; 0.25 M Tris base, 2.5 M NaCl, pH 7.36; stock solution).** A 5× TBS stock solution was prepared by dissolving 30.26 g of Trizma Base and 145 g of NaCl in 850 mL of deionized water with stirring until fully dissolved. The pH was adjusted to 7.36 using 37% HCl (15 mL were added initially, followed by gradual addition until pH 7.36 was reached). Deionized water was added if needed to reach the desired final volume. The solution was stored at 4 °C for several months.

**1× Tris Buffered Saline with Tween-20 (TBS-T; 0.05 M Tris base, 0.5 M NaCl, pH 7.36; 0.05% Tween-20; 1 L).** A 1× TBS-T solution was prepared by diluting 200 mL of 5× TBS stock solution with 800 mL of deionized water. Tween-20 was added to a final concentration of 0.05% (500 µL per 1 L), and the solution was mixed thoroughly until homogeneous. The buffer was stored at 4 °C for several months.

## 3 Methods

### 3.1. Establishment of patient-derived bile organoids

**CRITICAL STEP.** Once ERCP is completed, keep bile at 4 °C and process within 2 h of collection.

#### 3.1.1. Before you begin ◊ Timing 30 minutes (plus ≥ 1h plate pre-warming)

Bile must be processed fresh. Before starting the protocol:

- Pre-warm the culture plate where organoids will be seeded. Pre-warm for ≥ 1h at 37 °C in the incubator.
- Cool the centrifuge to 4 °C.
- Thaw Matrigel slowly on ice or at 4 °C. Slow thawing is critical to prevent bubble formation and premature gelation.
- Prepare CCM (prepare reasonable volumes and avoid storing complete medium for >1 week at 4 °C).

#### 3.1.2. Pre-culture processing of bile samples ◊ Timing ∼60–75 min

a) Bile (4–12 mL) collected at endoscopic facility in sterile additive-free tubes is distributed into 1mL aliquots and centrifuged for 10 min at 1,800 × g, 4 °C.
b) Remove the supernatant (bile can be kept for subsequent analyses) and resuspend the resulting pellet from each 1-mL aliquot in 1 mL of cold BCM.

**NOTE#2.** A visible pellet is often not observed. Proceed with the subsequent steps regardless, as organoids can still develop even without a discernible pellet.
**NOTE#3.** If a relatively large yellow–brown loose pellet is observed, dark aggregates may be observed when seeding the drop (step n). It is important to continue with the protocol, as in our hands, organoids have still been successfully generated from some samples under these conditions.
c) Filter all the BCM aliquots through a 70 µm nylon cell strainer placed over a conical 50 mL tube.
d) Centrifuge the conical 50ml tube at 400 x g for 5 min at 4 °C.
e) Discard the supernatant and wash the pellet: resuspend in 8 mL BCM and centrifuge at 400 × g for 5 min at 4 °C, repeat two additional times (total 3 washes).

#### 3.1.3. Erythrocyte lysis (if required) ◊ Timing 10–15 min

**CRITICAL STEP.** If a red pellet is observed after washing, lyse erythrocytes as follows:

f) Add 500 µL 1× ACK lysis buffer to the pellet and flick the tube to dislodge the pellet. Incubate for 2 min at 37 °C.
g) Stop the reaction by adding 2.5 mL of cold BCM (5× the ACK volume).
h) Centrifuge for 5 min at 400 × g, 4 °C and discard the supernatant.

* If the pellet remains clearly red, repeat the ACK lysis step (f-h).

**NOTE#4.** Longer or repeated lysis steps may compromise sample integrity; in our workflow, we avoid performing more than two consecutive ACK lysis cycles on the same sample; proceed even if a slight red tinge persists.

#### 3.1.4. Matrigel embedding and culture initiation ◊ Timing ∼45–60 min

i) Retrieve the pre-warmed untreated 24-well plate from the incubator and place it in the biosafety cabinet.
j) Take the tube from step h and ensure that the supernatant is completely removed. Add cold Matrigel according to the number of domes to be prepared (35 µL per dome) and resuspend the pellet rapidly but gently using pre-chilled, cut tips.

**CRITICAL STEP.** Keep Matrigel at 4 °C or on ice during handling; use single-use aliquots to avoid repeated freeze–thaw cycles.
**NOTE#5.** Typically, prepare one dome if no visible pellet is observed, or two domes if a visible pellet is obtained.
k) Dispense 35 µL domes of the Matrigel–cell suspension onto the center of a well of the pre-warmed plate.

**NOTE#6.** Volumes >35 µL may reduce dome integrity and roundness.
l) Immediately invert the plate and incubate for 35–40 min at 37 °C to allow polymerization.

**CRITICAL STEP.** Inverting the plate prevents cells from settling at the bottom of the plate and helps ensure uniform embedding within the Matrigel.
m) Carefully return the plate to the upright position and add 600 µL of pre-warmed CCM per well.
n) Maintain organoids at 37 °C in a humidified atmosphere with 5% CO₂.

### 3.2. Organoid maintenance and expansion

a. Inspect organoids under brightfield microscopy at least twice per week and document morphology (e.g., cystic vs filled/compact organoids, presence of darkening/debris).
b. Replace CCM twice per week using pre-warmed medium.

**NOTE#7.** Organoid outgrowth timing and establishment efficiency may vary across samples. In our hands, initial organoid outgrowth is typically observed within 2-5 days (**Figure 1A**).

**Figure 1.**
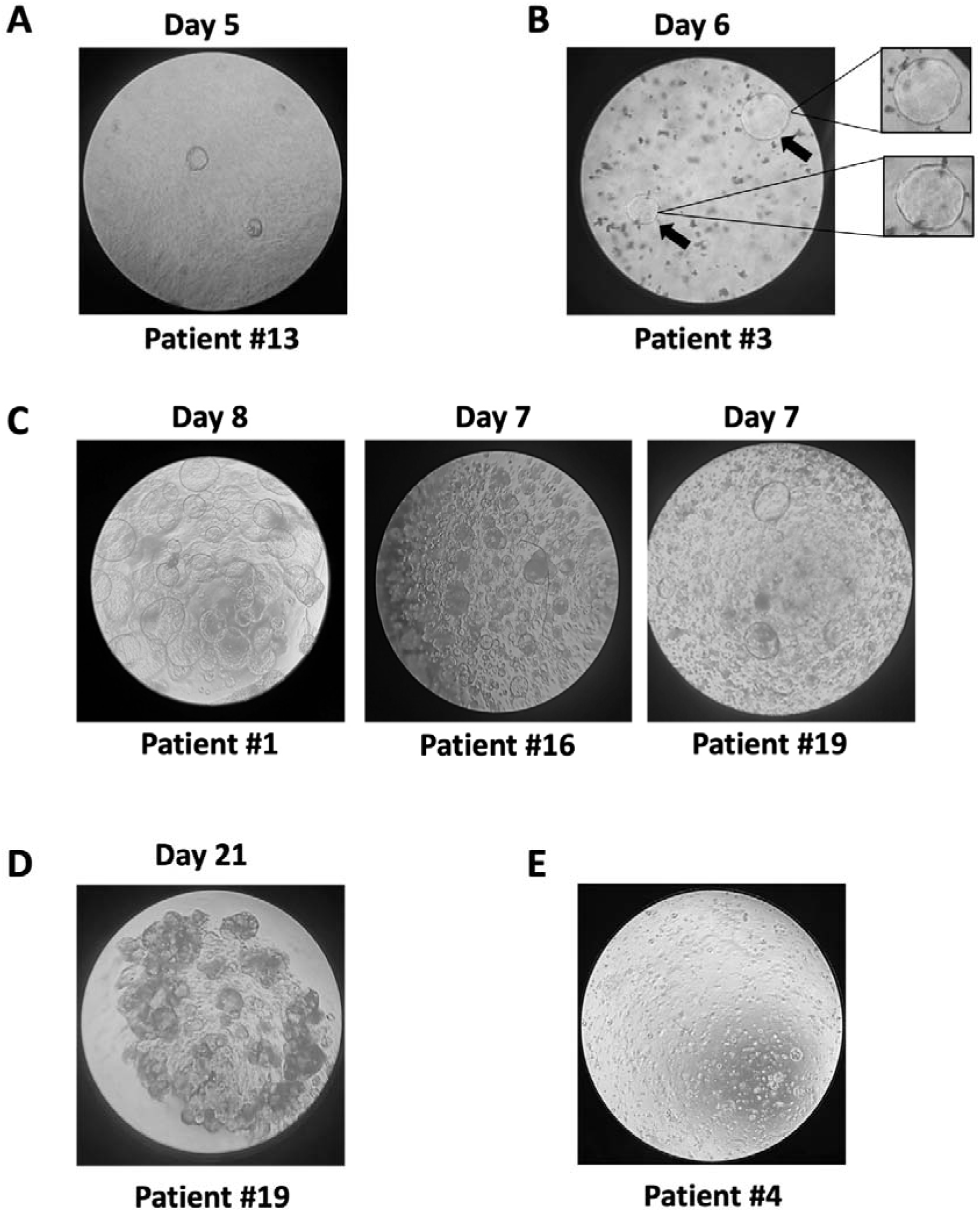
Representative brightfield microscopy of bile-derived organoid cultures. Brightfield microscopy at 4x magnification illustrates the developmental stages and culture quality of organoids derived from different patients. **(A)** Initial organoid growth observed at day 5 post-collection (Patient #13). **(B)** Organoid formation at day 6 (Patient #3) from a sample presenting a large yellow–brown loose pellet, with organoids emerging amidst dark aggregates. **(C)** Representative images from three distinct patients showing a high density of large, well-formed organoids at optimal confluence for passaging. **(D)** Morphological darkening of organoid structures, typically indicating reduced viability or the onset of cell death. **(E)** Established organoids 3 days post-passaging displaying exponential growth phase and readiness for cryopreservation.

**NOTE#8**. **Figure 1B** shows representative organoids at day 6 post-seeding from cultures initiated under the conditions described in NOTE#3 presenting debris or dark aggregates.

**NOTE#9.** After successful establishment, users may consider short “maturation windows” by reducing WNT3A/R-spondin stimulation and selected growth factors to favor a more differentiated cholangiocyte phenotype, while maintaining standard CCM for routine expansion and biobanking (Huch et al., 2015; Sampaziotis et al., 2017; Roos et al., 2021a).

### 3.3. Organoid passaging ◊ Timing ∼1.5 h

**NOTE#10.** Passage organoids when the Matrigel dome is densely populated with large, well-formed organoids (see **Figure 1C** for representative brightfield images). A typical split ratio is 1:3.

**NOTE#11.** Progressive darkening of organoids often indicates reduced viability and/or onset of cell death. In our experience, this typically reflects overgrowth and suggests that organoids should have been passaged earlier (see **Figure 1D** for representative brightfield images).

Remove CCM from the well that are going to be split. If multiple domes from the same patient/condition are splitted on the same day, we recommend to pool them and then generate the corresponding number of domes.

a) Remove CCM from wells to be passaged.
b) Add 500 µL of ice-cold BCM to the well and gently disrupt the Matrigel dome using circular pipetting movements. Repeat this step twice to maximize recovery. Transfer the suspension into a 15 mL conical tube containing 10 mL ice-cold BCM.

**NOTE#12**. If passaging multiple domes from the same patient/condition on the same day, pool all domes into the same 15 mL conical tube.
c) Centrifuge at 600 × g for 5 min at 4 °C. Carefully remove the supernatant without disturbing the pellet.
d) Enzymatic digestion (TrypLE). Add 500 µL to 1 mL TrypLE, depending on pellet size (adapted from (Driehuis et al., 2020)).

**NOTE#13**. As a guideline, 1 mL TrypLE is typically used to digest material from >3 Matrigel domes containing multiple organoids.
e) Gently tap the tube to detach organoids from the bottom. Incubate for 3 min at 37°C in a water bath.
f) Mechanically disrupt organoids by pipetting up and down 10 times.
g) Stop digestion by adding 5× the TrypLE volume of BCM.
h) Centrifuge at 300 × g for 5 min at 4 °C and discard the supernatant.
i) Resuspend the pellet in the corresponding volume of Matrigel (according to the number of domes to be prepared according to the original number of domes harvested; 35 µL per dome) and plate the domes onto a pre-warmed 24-wells plate (step j in 3.1.4.). Immediately invert the plate and incubate for 35-40 min at 37 °C to allow polymerization.
j) Return the plate to the upright position and add 600 µL of pre-warmed CCM per well.

### 3.4. Cryopreservation and Thawing of Organoids

The following protocols are adapted from the methodology described by (Driehuis et al., 2020).

### 3.4.1. Cryopreservation ◊ Timing ∼30–45 minutes

**CRITICAL STEP.** Organoids should be frozen 3–5 days post-passaging to ensure they are in exponential growth, this improves post-thaw survival and their ability to resume growth after thawing (see **Figure 1E** for representative brightfield images of organoids at the time of freezing).

**NOTE#14**. We typically freeze one Matrigel dome per cryovial. If multiple domes are harvested on the same day, they may be pooled and then aliquoted into as many cryovials as the number of domes originally collected.

a) Mechanically disrupt the Matrigel domes by adding 500 µL of ice-cold BCM per well. Pipette using gentle circular motions until the domes are fully detached and the organoids are evenly suspended.
b) Transfer the disrupted domes to a single pre-chilled 15 mL conical tube, pooling the material from different domes.
c) Centrifuge at 300 x g for 5 minutes at 4°C.
d) Carefully aspirate the supernatant without disturbing the organoid pellet.
e) Add 500 µL of Recovery™ Cell Culture Freezing Medium per Matrigel dome originally harvested.
f) Gently resuspend the pellet in the freezing medium and dispense into cryovials.
g) Place the cryovials immediately at -80 °C. After 3-4 days, transfer the vials to liquid nitrogen for long-term storage.

#### 3.4.2. Thawing ◊ Timing ∼1 hour

**CRITICAL STEP.** Rapid thawing is essential to maintain organoid viability. The transition from liquid nitrogen to 37 °C must be as fast as possible.

a) Retrieve the cryovial from storage and immediately place it in a 37 °C water bath. Hold the vial until only a small ice crystal remains.
b) Under sterile conditions perform gentle “up and down” pipetting to resuspend the organoids.
c) Transfer the suspension to a 15 mL conical tube and add an additional 4 mL of BCM.
d) Centrifuge at 300 x g for 5 minutes at 4 °C.
e) Aspirate the supernatant completely.
f) Resuspend the organoid pellet in the appropriate volume of Matrigel and seed as droplets (35 µL domes step j in 3.1.4.) onto a 37 °C pre-warmed culture plate.
g) Immediately invert the plate and incubate for 35–40 min at 37 °C to allow polymerization.
h) Carefully return the plate to the upright position and add 600 µL of pre-warmed CCM per well.
i) Maintain organoids at 37 °C in a humidified atmosphere with 5% CO₂.

### 3.5. Gene expression knockdown in bile-derived organoids

#### 3.5.1. Gene silencing in dissociated organoids ◊ Timing ∼7 h

**SCOPE.** Organoids are dissociated to a near single-cell suspension and gene expression knockdown in performed prior to re-embedding, enabling assessment of the effects of gene silencing on organoid initiation and formation.

**NOTE#15.** In this option, the silencing step is performed in 24-well plates (final volume 1 mL/well) under gentle rocking at 32 °C for 5 h.

a) Collect pre-formed organoids and dissociate them using the TrypLE-based digestion procedure described in Section 3.3 (Organoid passaging) steps, adapting volumes as needed to obtain a near single-cell suspension suitable for counting.

**CRITICAL STEP**. Efficient Matrigel removal and generation of a homogeneous suspension in a suitable volume for accurate cell counting are essential for reproducible seeding and gene silencing.

b) Resuspend the cells and count them using a Neubauer hemocytometer. Place 50,000 cells per condition in 800 µL Opti-MEM + 10% FBS in 24-well plates.

**NOTE#16.** To minimize variability, prepare a single master cell suspension in Opti-MEM + 10% FBS containing the total number of cells required for all wells plus a small excess to account for pipetting losses. For example, to seed 6 wells at 50,000 cells/well in 500 µL/well, prepare a master mix of 3.2 mL containing 320,000 cells (equivalent to 100,000 cells/mL) and dispense 500 µL per well. Mix gently but thoroughly before dispensing each aliquot to ensure homogeneous cell distribution.
c) Prepare siRNA transfection mixes using Lipofectamine RNAiMAX according to the manufacturer’s instructions. For each siRNA condition, prepare a single master siRNA–RNAiMAX mix in 200 µL Opti-MEM per well containing siRNA at 50 nM (plus ∼5–10% excess for pipetting losses). Add 200 µL of the corresponding siRNA–RNAiMAX mix to each well that already contains 800 µL of the cell suspension. This results in a final volume of 1 mL per well and a final siRNA concentration of 10 nM.
d) Incubate plates at 32 °C for 5 h under gentle agitation on a rocking platform placed inside the incubator.

**CRITICAL STEP**. During the incubation, cells should remain in suspension to maximize siRNA uptake and to prevent cell attachment. Use low-attachment plates, a relatively large working volume (1 mL/well) and gentle rocking throughout the incubation.
e) After 5 h, collect cells and centrifuge at 200 × g for 5 min at 4 °C. Carefully remove the supernatant without disturbing the cell pellet.
f) Re-embed the cell pellet in Matrigel and restart culture as described in Section 3.3 (Organoid passaging), steps h-i.

#### 3.5.2. Gene silencing in fully formed organoids ◊ Timing 7 hours

**SCOPE.** Pre-formed organoids are silenced while embedded/maintained as 3D structures, enabling assessment of the effects of gene silencing in established organoids (e.g., viability, growth, morphology and downstream molecular readouts).

**NOTE#17.** In this option, the silencing step is performed in 1.5 mL conical tubes (final volume 1 mL/tube) under gentle rocking at 32 °C for 5 h.

a) Without removing the culture medium from the well, carefully retrieve the Matrigel dome containing organoids from the culture well without disrupting its structure. Transfer it into a conical-bottom 1.5 mL tube containing 700 µL cold PBS. A small sterile spatula or spoon-shaped lifter can be used to handle the dome safely.

**NOTE#18.** Optionally, two domes from the same condition can be combined in one tube (do not exceed two). However, processing domes separately is recommended whenever possible, as it facilitates more uniform Matrigel disruption and handling, and may improve the overall silencing efficiency.
b) Centrifuge 1 min at 200 × g, 4 °C to settle the Matrigel dome without deforming it. Carefully remove PBS.

**CRITICAL STEP.** To avoid aspirating organoids, remove PBS using a 200 µL gel-loading tip.
c) Wash the organoids by adding 700 µL cold PBS and gently pipetting up and down four times using a cut P1000 tip to remove residual Matrigel while maintaining organoid integrity.
d) Centrifuge again (1 min at 200 × g, 4°C) and carefully remove all PBS.
e) Add 800 µL Opti-MEM + 10% FBS in two steps: first add 2 × 200 µL, gently detach organoids from residual Matrigel with 2–3 gentle up-and-down pipetting using a cut tip, then add the remaining 400 µL.

**CRITICAL STEP.** Avoid vigorous pipetting to preserve organoid structure.
f) Prepare siRNA transfection mixes using Lipofectamine RNAiMAX according to the manufacturer’s instructions. For each condition, prepare a siRNA–RNAiMAX mix in 200 µL Opti-MEM containing siRNA at 100 nM (plus ∼5–10% excess if preparing multiple tubes). Add 200 µL of this mix to each tube already containing 800 µL Opti-MEM + 10% FBS (step e). This yields a final volume of 1 mL per tube and a final siRNA concentration of 20 nM.
g) Incubate tubes at 32°C for 5 h under continuous gentle agitation on a rocking platform placed inside the incubator. Secure tubes to prevent tipping (e.g., place 1.5 mL tubes inside a 50 mL conical tube cushioned with paper towels).

**CRITICAL STEP.** Continuous gentle agitation improves exposure of organoids to the silencing mix and helps maintain uniform conditions within the tube.
h) After 5 h, centrifuge 1 min at 200 × g, 4 °C. Carefully discard the siRNA-containing medium without aspirating organoids.
g) Re-embed organoids in Matrigel and restart culture as described in Section 3.3 (Organoid passaging), steps h–i.

### 3.6. Immunohistochemistry of organoids ◊ Timing 4 days

a) Without removing the culture medium from the well, carefully retrieve the Matrigel dome containing organoids from the culture well without disrupting its structure. Transfer it into a conical-bottom 1.5 mL tube containing 200 µL cold PBS. A small sterile spatula or spoon-shaped lifter can be used to handle the dome safely.

**CRITICAL STEP.** Add PBS to the 1.5 mL tube before transferring organoids. The buffer facilitates organoid release from the spatula and prevents organoids from sticking to the plastic.
**NOTE#19**. Try to collect the entire droplet without disrupting it. If you recover most of it on the first attempt, avoid repeating the process, as attempting to collect it again may disrupt the organoids.
b) Adjust the volume to 1 mL with PBS and gently invert the tube 3–5 times to wash the sample.
c) Centrifuge for 1 min at 200 × g, 4 °C. Carefully remove the supernatant.

**NOTE#20.** To minimize the risk of aspirating organoids, remove the supernatant using a 200 µL gel-loading tip.
d) Add 1.2 mL of 4% paraformaldehyde (PFA), ensuring complete coverage of the sample. Fix for 1 h at room temperature: incubate the first 30 min under gentle rotation, then stop rotation and leave the tube upright for an additional 30 min at room temperature to allow organoids to settle at the bottom.

**NOTE#21.** To enable gentle and uniform rotation, place the 1.5 mL tube inside a 50 mL conical tube and stabilize it (e.g., between two layers of paper). Mount the 50 mL tube on a roller/rotator designed for conical tubes.
e) Carefully remove the PFA solution.
f) Wash organoids (×3). Add 700 µL PBS, gently invert to resuspend, centrifuge for 2 min at 200 × g, 4 °C, and remove the supernatant as in step (c). Repeat for a total of three washes.

**PAUSE POINT.** After the three PBS washes (step f), fixed organoids can be kept in PBS at 4 °C for up to 72 h before proceeding to agarose pre-embedding and paraffin processing. Although we recommend completing the workflow without interruption when possible, this pause point enables flexible scheduling.
g) Prepare a 2% agarose solution by heating it to boiling in a microwave oven (or by melting a prepared stock solution).
h) Centrifuge for 5 min at 200 × g and remove the supernatant. Place the 1.5 mL tube in a water bath at 55–60 °C.

**NOTE#22.** Keeping the tube in the hot water bath prevents the agarose from solidifying prematurely.
i) Gently add 200 µL of 2% agarose using a cut P1000 tip and mix the organoids carefully.
j) Gently, but rapidly, transfer the organoids/agarose mixture into the cryomold.

**NOTE#23.** Placing the cryomolds on a dark surface helps visualize the transfer of organoids from the tube to the cryomold.
k) Keep the cryomolds at 4 °C until the agarose is completely solidified (∼20 minutes).
l) Carefully remove the solidified agarose containing the embedded organoids using a new razor blade and place it into a pre-labeled tissue cassette.
m) Transfer the tissue cassette to 70% ethanol until subsequent paraffin embedding process.
n) Proceed with the paraffin embedding workflow following a standard protocol for formaldehyde-fixed, paraffin-embedded (FFPE) samples.
o) Carefully trim the paraffin block and cut 3 µm sections using a rotary microtome. Float the ribbon of sections on a 42 °C water bath and collect them onto adhesive microscope slides (TOMO slides were used). Keep the slides in a vertical position until completely dry (overnight). Dried sections can then be used for hematoxylin–eosin (H&E) staining and immunohistochemistry.

**NOTE#24.** Organoids are often difficult to visualize by the naked eye in the sections; therefore, their presence should be confirmed using a light microscope. It is advisable to gently mark the position of the sections on the back of the slide with a diamond pencil to facilitate subsequent localization.
p) Bake slides in a 60°C drying oven for 30 minutes. Deparaffinized and rehydrated sections to running tap water.
q) Perform antigen retrieval using TE buffer (pH 9) in a Pascal Pressure Chamber at 95°C for 30 min. Slides were allowed to cool for 25 min at room temperature.
r) Block endogenous peroxidase by incubating the slides in 3% H₂O₂ in distilled water for 12 min at room temperature and briefly rinse in running tap water.
s) Wash slides in TBS-T for 5 min.
t) Gently dry the slides surrounding the tissue sections and add the primary antibodies, anti-EpCAM (Abcam, ab223582; 1:1000) or anti-CK19 (Abcam, ab52625; 1:200).
u) Incubate overnight at 4 °C in a humidified chamber.
v) Wash slides in TBS-T for 5 min.
w) Dry slides around sections and incubated with peroxidase labelled polymer conjugated goat anti-rabbit (EnVision+). Incubate for 30 min at room temperature.
x) After washing in TBS-T for 5 min, apply DAB+ solution for 10 min. Rinse in running tap water for 5 min.
y) Finally, lightly counterstain the sections with Harris hematoxylin, followed by dehydration and mounting.

### 3.7. Protein extraction and Western blot analysis ◊ Timing ∼36 h

a) Collect organoids from Matrigel and obtain a cell suspension as described in Section 3.3 (Organoid passaging), steps a–g, adapting volumes as needed.
b) Lyse cells in ice-cold RIPA buffer (see Reagent setup) supplemented with protease and phosphatase inhibitors for 20 min at 4 °C under constant rotation.

**CRITICAL STEP.** Keep samples and buffers cold to preserve protein integrity and phosphorylation states.
c) Sonicate lysates and centrifuge at 11,000 g for 20 min at 4 °C to remove insoluble material. Transfer the clarified supernatant to a new tube.
d) Measure protein concentration using the BCA assay (Pierce) according to the manufacturer’s instructions.
e) Perform Western blot analysis as previously described (Jiménez et al., 2019).

### 3.8. Total DNA isolation and molecular analyses

#### 3.8.1. Total DNA isolation ◊ Timing ∼1–2 h

a) Collect organoids and obtain a cell pellet as described in Section 3.3 (Organoid passaging, steps a-g), stopping after the final centrifugation step (prior to Matrigel re-embedding).
b) Extract total DNA using the automated Maxwell RSC Instrument with the Cultured Cells DNA kit (Promega, Madison, WI, USA) following the manufacturer’s instructions.
c) Measure DNA concentration using a NanoDrop spectrophotometer and/or Qubit, as appropriate for downstream applications.

#### 3.8.2. Next Generation Sequencing DNA analyses

Targeted sequencing was performed using 50 ng of DNA with the Oncomine™ Pan-Cancer Cell-Free Assay (Thermo Fisher Scientific, Waltham, MA, USA) according to the manufacturer’s instructions and as previously described (Arechederra et al., 2022, 2025).

#### 3.8.3. ULP-WGS for copy number alterations (CNAs)

Ultra–low-pass whole-genome sequencing (ULP-WGS) was performed using 5 ng of DNA as previously described (Sogbe et al., 2023) and according to the manufacturer’s instructions, where applicable. Copy-number profiling and tumour fraction (TF) estimation were conducted using the ichorCNA framework, following the recommended workflow for cfDNA to infer large-scale CNAs and aneuploidy patterns (Adalsteinsson et al., 2017).

## 4 (Anticipated) Results

### 4.1 Cohort description and organoid establishment rate

Bile samples were collected from 21 individuals undergoing ERCP for biliary strictures (**Table 1**). The cohort comprised 5 cases with benign biliary stenosis due to choledocholithiasis and 16 cases with a malignant cause of stricture, including 13 CCAs, 1 gallbladder adenocarcinoma, and 2 ampullary tumors.

Successful establishment was defined as the generation of organoids that could be expanded and passaged at least once. Using the workflow described in Section 3.1, bile-derived organoids were successfully established in 17/21 samples (81%) (**Table 2**). Establishment rates differed by diagnosis (**Table 2**), with success rates of 85% (11/13) for CCA, 100% (2/2) for ampullary tumor, 100% (1/1) for GBC, and 60% (3/5) for choledocholithiasis.

**Table 2.** Establishment success rate of bile-derived organoids according to diagnosis. The study cohort (n=21) consisted of 5 cases of benign stenosis (choledocholithiasis) and 16 malignant cases, including 13 CCA, 1 GBC, and 2 ampullary tumors. Percentage of establishment success per diagnostic group, defined as cultures successfully reaching at least one passage.

|  | Established | Non-established | Total | Success rate (%) |
| --- | --- | --- | --- | --- |
| **Choledocholithiasis** | 3 | 2 | 5 | 60 |
| **Cholangiocarcinoma** | 11 | 2 | 13 | 85 |
| **Gallbladder cancer** | 1 | 0 | 1 | 100 |
| **Ampullary tumor** | 2 | 0 | 2 | 100 |

### 4.2 Morphology, outgrowth kinetics and routine expansion

Of the successfully established cultures, organoids from 7 patients (2 choledocholithiasis, 4 CCA and 1 GBC) were expanded and characterized in detail in this study. Brightfield microscopy revealed the appearance of epithelial structures compatible with biliary organoid outgrowth within ∼2–3 weeks after seeding. The time to first detectable organoid outgrowth varied across samples (range [4–19] days; median [11.71] days), consistent with inter-individual variability in bile cellular content and sample quality (examples in **Table 3**).

**Table 3.** Time to establishment of bile-derived organoids. Time to establishment (days) for the subset of 7 patients (2 choledocholithiasis, 4 CCA, and 1 GBC) characterized in detail in this study.

| Patient ID | Diagnosis | Time to establishment (days) |
| --- | --- | --- |
| #1 | Choledocholithiasis | 4 |
| #18 | Choledocholithiasis | 19 |
| #13 | CCA | 16 |
| #14 | CCA | 12 |
| #15 | CCA | 10 |
| #19 | CCA | 10 |
| #16 | GBC | 11 |

In the choledocholithiasis-derived cultures, organoids displayed a typical cystic morphology, forming a single layer of epithelial cells positive for CK19 and EpCAM surrounding a central lumen (**Figure 2A)**. Across samples from individuals with malignant strictures, we observed marked intra- and inter-sample heterogeneity in organoid morphology. In most cases, cultures initially contained mixed organoid populations of normal cholangiocyte-like and tumoral-like organoids (**Figure 2B-C**). Representative brightfield images and H&E, CDK19 and EPCAM immunohistochemistry of the organoids obtained from a CCA patient (**Figure 2B**) and a GBC patient (**Figure 2C**) are shown. In the H&E staining examples of organoids with normal (1) and tumoral (2) morphology are shown at higher magnification.

**Figure 2.**
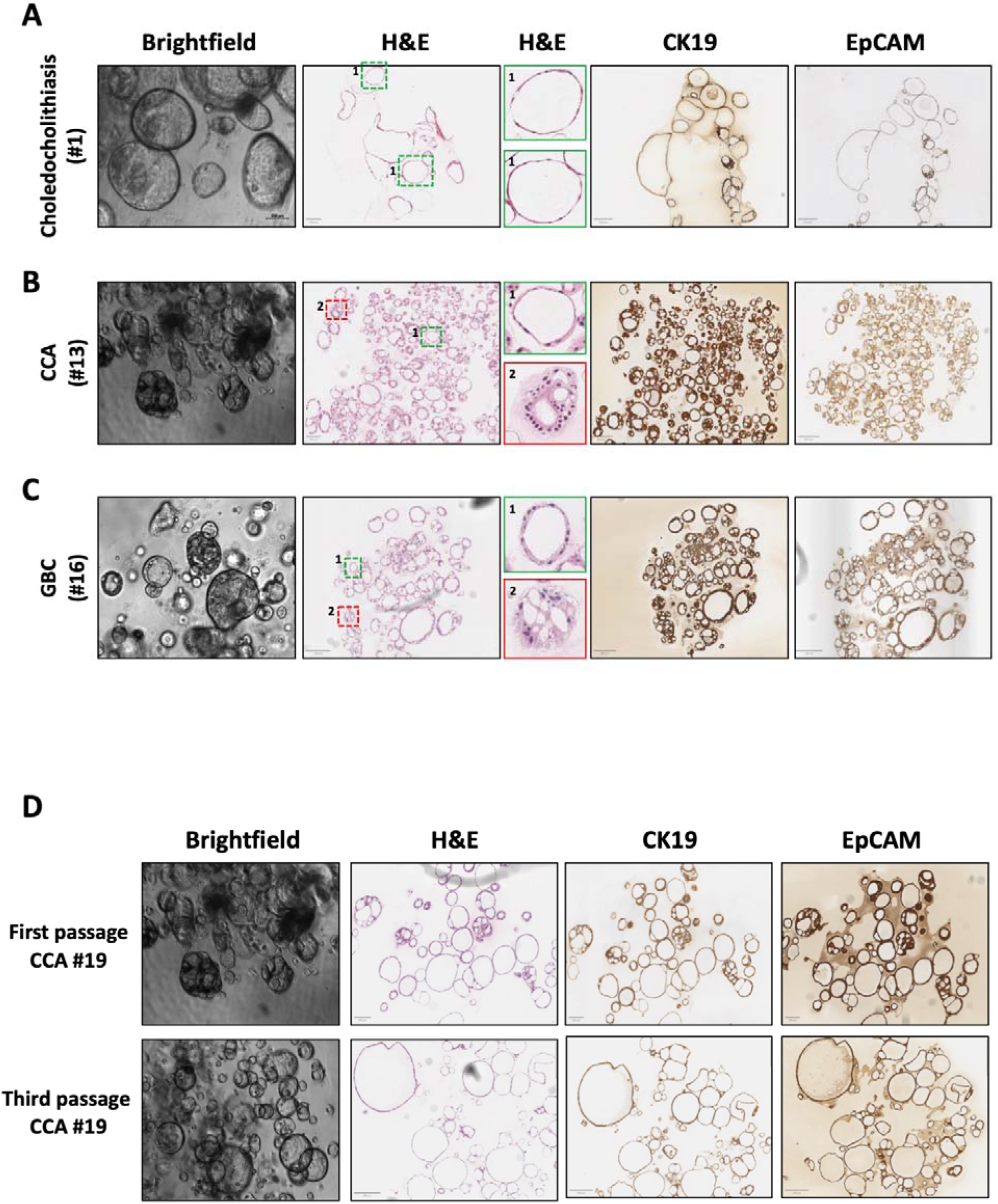
Morphology and histological characterization of bile-derived biliary organoids and culture drift during expansion. **(A)** Representative brightfield images and corresponding H&E, CK19 and EpCAM immunohistochemistry from a choledocholithiasis-derived culture (Patient #1), showing typical cystic cholangiocyte-like organoids. **(B–C)** Representative brightfield images and corresponding H&E, CK19 and EpCAM immunohistochemistry from one CCA-derived culture (Patient #13) (B) and one GBC-derived culture (Patient #16) (C), illustrating the coexistence of normal cholangiocyte-like and more tumour-like organoid populations within the same culture. Insets show magnified views of representative organoids: normal cholangiocyte-like organoids (1) are outlined in green and tumour-like organoids (2) in red; note the larger size and increased structural complexity of the tumour-like structures. **(D)** Example of phenotypic drift during routine passaging of a CCA-derived culture (Patient #19), illustrating a shift toward a predominantly cystic, cholangiocyte-like morphology over expansion (top: first passage; bottom: third passage).

However, under routine expansion and passaging, the cultures often drifted over time toward an almost exclusive cystic phenotype, consistent with the preferential outgrowth of non-malignant cholangiocyte-derived organoids (**Figure 2D**). Therefore, physical separation by manual handpicking, as previously described (Broutier et al., 2017), would be required to isolate and maintain the tumor-derived organoid population.

### 4.3 Pilot targeted NGS and ULP-WGS in selected paired bile–organoid samples

To evaluate whether bile-derived organoids capture the genomic features detected in the matched bile samples, we performed targeted NGS using the Oncomine™ Pan-Cancer Cell-Free Assay in selected paired bile–organoid samples (**Table 4**). In a choledocholithiasis control (patient #1, the same *TP53* polymorphic variant (p.R213= consistent with a germline polymorphism) was detected with comparable MAF values in bile and the matched organoid culture (52,21% and 47.32%, respectively). In a representative CCA case (patient #13), bile cfDNA sequencing detected again the *TP53* p.R213= variant but together with mutations at low-MAF in *EGFR* (R451C; 0.08%) and *KRAS* (G12D; 0.02%). In the matched organoid culture, *TP53* p.R213= was also observed and *KRAS* (G12D; 0.12%) was detected, whereas *EGFR* (R451C) was not detected. In a representative GBC case (patient #16), bile cfDNA sequencing identified a low-allele-frequency *ERBB3* mutation (D297Y; 0.015%). The matched organoid culture harbored the same variant at a higher MAF (D297Y; 1.11%). Overall, these results indicate that targeted sequencing is feasible in organoid-derived DNA and can recover bile-detected alterations.

**Table 4.** Targeted NGS in selected paired bile–organoid samples. List of mutations identified in the paired bile cfDNA and bile-derived organoid gDNA using the Oncomine™ Pan-Cancer Cell-Free Assay. Gene, amino acid change and percentage of mutant-allele frequency are indicated. Abbreviations: cfDNA: cell-free DNA; gDNA: genomic DNA.

| Patient ID | Diagnosis | NGS mutational Analysis (PanCancer) |  |
| --- | --- | --- | --- |
|  |  | Bile | Bile-derived organoids |
| #1 | Choledocholithiasis | *TP53 polymorphism* (R213=; 52,2141%) | *TP53 polymorphism* (R213=; 47,3203%) |
| #13 | CCA | *KRAS* (G12D; 0.02%); *EGFR* (R451C; 0.0788%); *TP53 polymorphism* (R213=; 45,5474%) | *KRAS* (G12D; 0,1272%); *TP53 polymorphism* (R213=, 49.8686%) |
| #16 | GBC | *ERBB3* (D297Y; 0.015%) | *ERBB3* (D297Y; 1.1178%) |

In addition, we assessed genome-wide copy number alterations by ULP-WGS in selected paired samples (**Table 5**). In the choledocholithiasis control (patient #1), TF estimates were 0 in both bile and organoid-derived DNA, as expected for a non-malignant condition. In contrast, in a CCA case (patient #14), ULP-WGS revealed a detectable TF in bile of 39% and a lower TF of 8.4% in the corresponding organoid culture, supporting the feasibility of CNA profiling from both bile and bile-derived organoids and indicating that organoid-derived DNA can retain tumor-associated signals detected in bile.

**Table 5.** ULP-WGS in selected paired bile–organoid samples. Percentage of tumor fraction detected by ULP-WGS analyses by ichorCNA in the paired bile cfDNA and bile-derived organoid gDNA. Abbreviations: cfDNA: cell-free DNA; gDNA: genomic DNA.

| Patient ID | Diagnosis | ULP-WGS |  |
| --- | --- | --- | --- |
|  |  | Bile | Bile-derived organoids |
| #1 | Choledocholithiasis | 0 | 0 |
| #14 | CCA | 39% | 8,40% |

### 4.4. siRNA gene expression knockdown in dissociated and formed organoids

We next evaluated the feasibility of siRNA-mediated gene silencing using two complementary workflows: silencing in dissociated organoids prior to re-embedding (Section 3.5.1) and silencing in fully formed organoids (Section 3.5.2).

In the dissociated workflow we followed the protocol described in (Aloia et al., 2019). Pre-formed organoids (**Figure 3A**) were dissociated to a single-cell–enriched suspension, transfected with siRNAs, and re-embedded in Matrigel (**Figure 3A**). As a proof of concept, we targeted X mRNA using siX, with siGL as a negative control. After re-embedding, parallel wells were processed at defined time points for knockdown validation (96 h), while replicate wells were maintained for long-term monitoring of organoid re-formation and outgrowth (13 days). At 96 h post-transfection, the X protein levels were assessed by Western blot and were found reduced compared with control conditions (**Figure 3B**). Although at different density, newly formed organoids were observed by day 13 in both the siGL control and the siX condition, indicating preservation of organoid-forming capacity following this procedure (**Figure 3A**).

**Figure 3.**
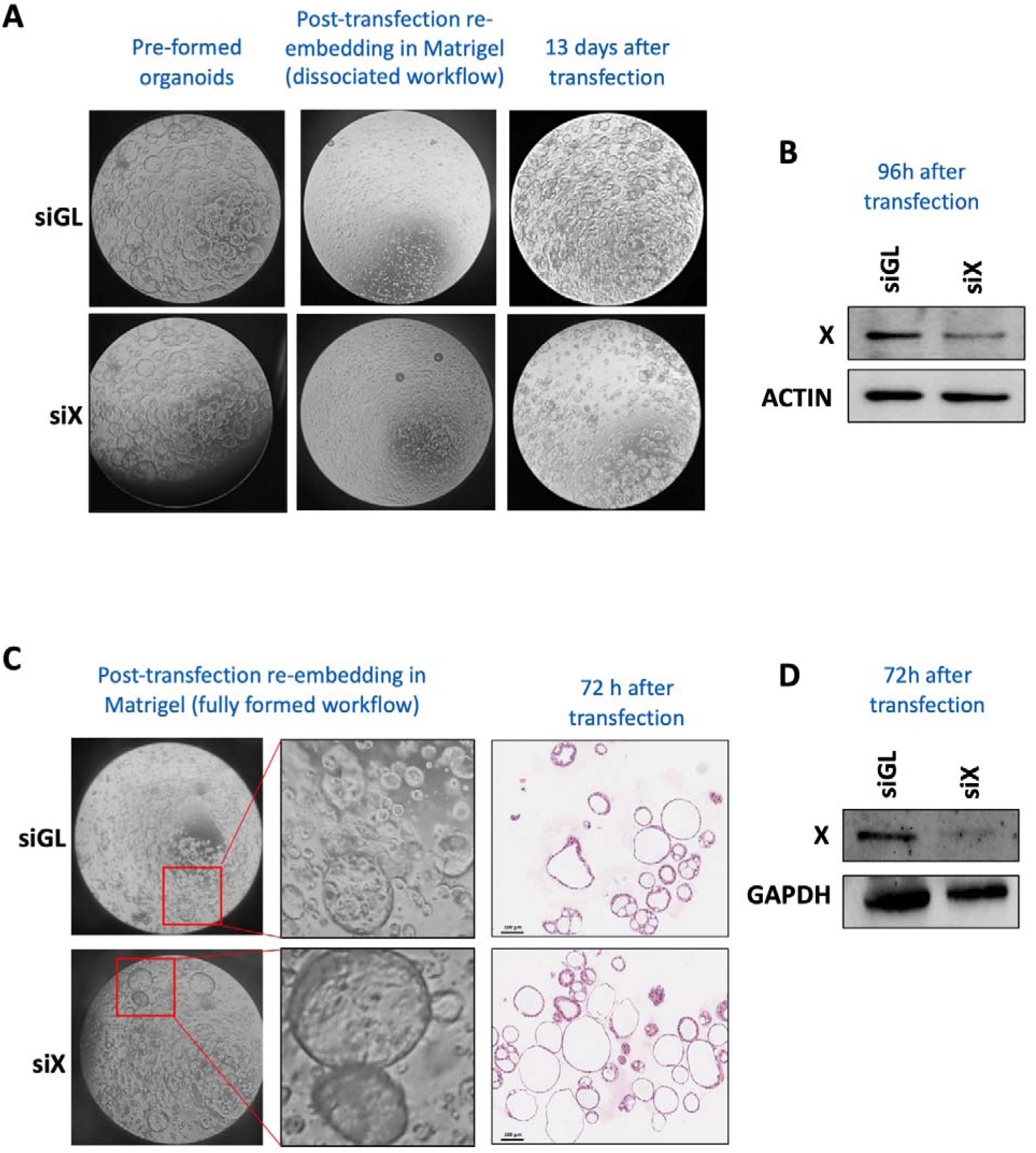
Evaluation of siRNA-mediated silencing in dissociated and fully formed bile-derived organoids. Representative brightfield microscopy and protein expression analysis demonstrate knockdown efficiency across two transfection workflows. **(A)** Brightfield images of the dissociated organoid silencing workflow, showing pre-formed organoids, post-transfection re-embedding in Matrigel, and organoid reconstitution 13 days after transfection with either siRNA targeting the gene of interest (siX) or a non-targeting control (siGL). **(B)** Western blot analysis of protein X levels 96 hours post-transfection in the dissociated workflow, with actin as a loading control. **(C)** Brightfield and immunohistochemical analysis of the fully formed organoid silencing workflow 72 hours post-transfection with siRNA targeting the gene of interest (siX) or a non-targeting control (siGL). **(D)** Western blot analysis of protein X levels 72 hours post-transfection in the fully formed workflow, with GAPDH as a loading control.

In the fully formed organoid workflow, pre-formed organoids (**Figure 3C**) were transfected with siX, using siGL as a control. At 72 h after transfection, histological assessment (H&E) supported maintenance of organoid structural integrity following the silencing procedure (**Figure 3C**) as well as knockdown efficiency by Western blot (**Figure 3D**), showing reduced X protein levels compared with control siRNA conditions (siGL).

### 4.5. Pitfalls, limitations and troubleshooting

This protocol enables the generation and downstream manipulation of bile-derived organoids; however, several factors can affect establishment efficiency, culture behavior and assay performance. Common pitfalls and recommended troubleshooting measures are summarized in **Table S2**.

## 5 Discussion

Robust human experimental systems are essential to move from descriptive studies toward actionable biology and translational testing. In the biliary tract, model availability has lagged behind other tissues, largely because access to healthy cholangiocytes is limited and tissue sampling is invasive, scarce, and often biased toward late-stage or surgically resected material. Tissue-derived cholangiocyte organoids were enabled by long-term expansion of adult human liver EpCAM+ ductal cells in organoid culture conditions (Huch et al., 2015) and further extended to extrahepatic/intrahepatic cholangiocyte organoid systems (Sampaziotis et al., 2017; Tysoe et al., 2019; Verstegen et al., 2020; Roos et al., 2022). In parallel, patient-derived organoids have been generated from biliary tract tumors, including cholangiocarcinoma and related BTC entities, providing clinically relevant platforms for modeling and functional testing (Broutier et al., 2017; Saito et al., 2019; Maier et al., 2021; Lee et al., 2023). While these tissue-based platforms have provided a major advance, their broad implementation is still constrained by sample accessibility and variability in tissue quality. In this setting, bile offers a practical alternative starting material that can be obtained during routine clinical procedures and, when processed appropriately, can yield expandable epithelial cultures suitable for standardized workflows. while preserving the limited histological tissue available in this tumor type (Soroka et al., 2019a, 2019b; Roos et al., 2021b; Kinoshita et al., 2023).

In line with recent community efforts to harmonize hepatopancreatobiliary organoid definitions and reporting standards (Marsee et al., 2021), we present an operator-oriented, step-by-step protocol for establishing biliary organoids from ERCP bile. Our workflow was developed and tested in a real-world cohort encompassing both benign and malignant strictures (5 benign biliary obstructions due to choledocholithiasis and 16 malignant strictures, including 13 CCA, 1 GBC, and 2 ampullary tumors). Beyond standard methods descriptions, we explicitly detail practical steps and decision points that are often under-reported yet can substantially affect success rates. By formalizing these critical control points, the protocol aims to improve reproducibility and facilitate implementation of bile as an input material for biliary organoid generation, complementing previously published bile-based protocols (Soroka et al., 2019b). Although ERCP is currently the most common source of bile for these applications and was used here, bile-derived organoids have also been initiated from alternative clinical routes (e.g., gallbladder-derived bile or percutaneous procedures), supporting broader applicability beyond ERCP-only workflows (Roos et al., 2021b).

A relevant consideration specific to bile from malignant strictures is the frequent coexistence of normal-like and tumor-like epithelial populations during early culture, reflecting the biological mixture of shed epithelial cells present in bile (Kinoshita et al., 2023). Although this heterogeneity can complicate interpretation when tumor-specific readouts are required, it can also be leveraged as a unique strength of bile-based sampling, as a single clinical specimen may provide access to both malignant and non-malignant organoids from the same individual, enabling within-patient comparisons and matched control systems. Practically, when the goal is to maintain both populations, physical separation by manual handpicking can be used to derive and propagate distinct lines from morphologically different organoids, as described in tumor organoid workflows (Broutier et al., 2017). Conversely, when the experimental goal is only to enrich for tumor-derived organoids, additional strategies may counteract the preferential outgrowth of normal cholangiocyte organoids under routine expansion conditions. In this context, Kinoshita and colleagues proposed complementary approaches including repeated passaging, xenografting, and selective pressure using an MDM2 inhibitor in TP53 mutation–harboring cases (Kinoshita et al., 2023). In line with this, our pilot targeted sequencing and ULP-WGS analyses in selected paired bile–organoid samples support that organoid-derived DNA can recover alterations detected in bile, although tumor-associated signals may be reduced compared with the matched bile specimen. Future applications of this workflow may benefit from systematic handpicking-based separation followed by molecular profiling, permitting the generation of paired normal-like and tumor-enriched lines from the same bile specimen for lineage-resolved comparisons.

Finally, to extend bile-derived organoids from descriptive modeling to functional interrogation, we integrate two complementary siRNA delivery workflows: one performed in dissociated organoid cells prior to re-embedding, and one performed directly in fully formed 3D organoids. Efficient gene perturbation in 3D cultures remains more variable than in 2D systems due to extracellular matrix barriers and the compact architecture of organoids. Accordingly, workflows frequently implement delivery of siRNAs at the dissociated/single-cell stage prior to re-embedding (Aloia et al., 2019), whereas knockdown in intact 3D structures is generally more challenging and less standardized (Laperrousaz et al., 2018; Morgan et al., 2018). By detailing handling conditions that preserve organoid integrity while enabling uptake and providing proof-of-concept knockdown validation (siRNA targeting gene X mRNA in both dissociated and fully formed organoids), this workflow supports functional gene interrogation in a clinically accessible biliary organoid system.

Overall, this protocol expands the practical repertoire of biliary models by enabling reproducible generation of bile-derived organoids across benign and malignant strictures and by incorporating functional siRNA perturbation approaches suitable for both early and established 3D cultures. We anticipate that standardized bile-derived organoid workflows will facilitate mechanistic studies in cholangiopathies, enable matched patient-derived comparisons, and support translational pipelines such as biobanking and functional testing.

## 7 Conflict of Interest

Authors declare that the research was conducted in the absence of any commercial or financial relationships that could be construed as a potential conflict of interest.

## 8 Author Contributions

CR: Methodology, Investigation, Visualization, Writing – original draft; JJV: Resources; LG: Methodology; AA-G: Resources; VJ-I: Resources; JC-G: Resources; MR: Resources; JR: Resources; MGF-B: Methodology; MH: Methodology; JU: Resources; MAA: Conceptualization, Funding

acquisition; CB: Conceptualization, Methodology, Investigation, Funding acquisition, Supervision, Writing – original draft; MA: Conceptualization, Methodology, Investigation, Visualization, Funding acquisition, Supervision, Writing – original draft. All authors have read and approved the final manuscript.

## 9 Funding

This work was supported by grants PI25/01079 and PI22/00471 funded by Instituto de Salud Carlos III (ISCIII) and co-funded by the European Union and INVES223049AREC fellowship from AECC to MA; Grant from Ministerio de Ciencia Innovación y Universidades MICINN-Agencia Estatal de Investigación integrado en el Plan Estatal de Investigación Científica y Técnica y Innovación, cofinanciado con Fondos FEDER PID2022-136616OB-I00/AEI/10.13039/501100011033 to MAÁ, CIBERehd; FIMA predoctoral and AECC predoctoral 2025 fellowships (PRDNA258508ROJO) to CR; European Union Horizon 2020 Transcan project [2022–784-024] to MAÁ and CB; Thermo Fisher 2022 Oncomine Clinical Research Grant to MAÁ and CB; AECT Eurorregión Nueva Aquitania Euskadi Navarra “Innovación Eurorregional” [2020/101] and [2023/2] to MAÁ and CB; Rolf M Schwiete Foundation grant 2024–040 to MAÁ and CB. Departamento de Salud Gobierno de Navarra [42/2021] to JU. This study is based upon work from COST Action Precision-BTC-Network CA22125, supported by COST. The generous support of Mr. Eduardo Avila is acknowledged.

## Acknowledgments

We particularly acknowledge the patients for their participation. We thank Dr. Pau Sancho-Bru and his team at the Institut d’Investigacions Biomèdiques August Pi i Sunyer (IDIBAPS) for their valuable advice. We also thank Luchi (Endoscopy Unit, Department of Gastroenterology and Hepatology, Navarra University Hospital) for her continuous assistance with sample collection. Finally, we acknowledge the technical support of the Imaging Core facility at CIMA (University of Navarra).

## Supplementary Material

**Table S1.** Composition of the conditioned culture medium (CCM) The table details the reagents, suppliers, stock concentrations, final working concentrations, and volumes required to prepare 10 mL of complete conditioned medium (CCM). The basal culture medium (BCM) was prepared as described in Section 2.5 and subsequently supplemented with Wnt3A-conditioned medium, growth factors, small-molecule inhibitors, and supplements. All components were added under sterile conditions using the indicated stock solutions. Single-use aliquots were employed to avoid repeated freeze–thaw cycles.

| Reagent | Reference | Manufacturer | Stock concentration | Final concentration | Volume for 10 mL |
| --- | --- | --- | --- | --- | --- |
| Basal Culture Medium (BCM) | 12634-010 | Gibco | – | – | 4.4 mL |
| Primocin | ant-pm2 | InvivoGen | 50 mg/mL | 100 µg/mL | 20 µL |
| 3dGRO™ L-WNT Conditioned Media | SCM105 | Merck | 2× | 1× | 5 mL |
| N2 supplement | 17502048 | Life Technologies | 100× | 1× | 100 µL |
| B27 supplement (w/o vitamin A) | 12587010 | Life Technologies | 50× | 1× | 200 µL |
| Nicotinamide | N0636-100G | Merck | 1 M | 10 mM | 100 µL |
| N-Acetylcysteine | A8199-10G | Sigma-Aldrich | 500 mM | 1.25 mM | 25 µL |
| Y-27632 (ROCK inhibitor) | Y-27632 | Tocris | 10 mM | 10 µM | 10 µL |
| [Leu15]-Gastrin I human | G9145-0.1MG | Sigma-Aldrich | 100 µM | 10 nM | 1 µL |
| Recombinant human HGF | 100-39H | PeproTech | 25 µg/mL | 25 ng/mL | 10 µL |
| Recombinant human FGF10 | 100-26 | PeproTech | 100 µg/mL | 100 ng/mL | 10 µL |
| Recombinant human EGF | AF-100-15 | PeproTech | 50 µg/mL | 50 ng/mL | 10 µL |
| A83-01 (TGF-β inhibitor) | SML0788 | Merck | 500 µM | 5 µM | 100 µL |
| Forskolin | 344270 | Calbiochem | 8 mM | 10 µM | 12.5 µL |

**Table S2.** Troubleshooting guide for the bile-derived organoid protocol. The table summarizes technical hurdles, their potential causes, and recommended corrective actions throughout the organoid workflow.

| Troubleshooting | Possible cause | Recommendation |
| --- | --- | --- |
| Reduced establishment success | Delayed processing; suboptimal transport conditions; sample-to-sample variability | Keep bile at 4 °C and process within 2 h. Use sterile additive-free tubes and exclude samples collected after contrast injection. |
| No visible pellet after centrifugation | Low cellular content in bile | Proceed with the protocol; organoids may still establish even in the absence of a visible pellet. Minimize sample loss during supernatant removal. |
| Prominent red pellet / erythrocyte contamination | Blood contamination during sampling | Perform ACK lysis, but avoid >2 consecutive cycles to prevent sample damage. Proceed if a slight red tinge persists after two cycles. |
| Matrigel dome spreads/ flattens or shows poor integrity | Matrigel warmed during handling; excessive dome volume; slow handling | Keep Matrigel on ice/at 4 °C, use pre-chilled plasticware and cut tips, and limit dome volume to 35 µL. Pre-warm plates ≥1 h at 37 °C. |
| Cells settle to the bottom of the dome during polymerization | Plate not inverted; delayed inversion | Invert the plate immediately after plating domes and polymerize at 37 °C for 35–40 min. |
| Organoids darken or show increased debris | Overgrowth, nutrient depletion, stress, suboptimal passaging timing | Refresh medium twice per week and passage organoids when well-formed and/or early darkening is observed. Avoid prolonged enzymatic digestion. |
| Poor recovery after passaging | Incomplete Matrigel disruption; over- or under-digestion | Ensure efficient dome disruption and use TrypLE volumes proportional to pellet size. Limit digestion time and standardize mechanical disruption (e.g., 10 pipettings). |
| Low silencing efficiency in dissociated workflow | Cells attach during incubation; uneven cell seeding; suboptimal siRNA/LNP ratios | Maintain cells in suspension during the 5 h incubation (use low-attachment formats where available, 1 mL/well total volume, and gentle rocking at 32 °C). Prepare a single master cell suspension and master transfection mixes per siRNA condition. |
| Loss of organoid integrity in formed-organoid workflow | Excessive pipetting or mechanical stress | Use cut tips and gentle handling during washes and media exchanges. Avoid vigorous mixing and minimize handling time outside the incubator. |
| Variable knockdown readout | Timing differences; protein vs transcript kinetics; line-to-line variability | Define consistent readout time points (e.g., 72–96 h) and confirm silencing by orthogonal methods (WB and/or qPCR). Optimize siRNA dose if needed. |
| Low DNA yield or inconsistent DNA quantification | Sample loss; inhibitors; measurement bias | Use automated extraction (as specified) and quantify DNA using Qubit and/or NanoDrop; interpret purity ratios |
|  |  | (A260/280 and A260/230). Consider re-extraction if purity is poor. |
| Discordant variants between bile and matched organoids | Tumor heterogeneity; sampling differences; selection during culture | Interpret bile and organoid profiling as complementary readouts. Report shared and private variants transparently and avoid over-interpreting discordance as technical failure. |
| Contamination (bacterial/fungal/mycoplasma) | Non-sterile handling; contaminated reagents | Use strict aseptic technique, test routinely for mycoplasma, and discard contaminated cultures. Antibiotics do not replace sterile workflow. |

## Notes

### Competing Interest Statement

The authors have declared no competing interest.

